# Effectors of the Spindle Assembly Checkpoint but not the Mitotic Exit Network are Confined within the Nucleus of Saccharomyces Cerevisiae

**DOI:** 10.1101/340091

**Authors:** Lydia R. Heasley, Jennifer G. DeLuca, Steven M. Markus

## Abstract

The Spindle Assembly Checkpoint (SAC) prevents erroneous chromosome segregation by delaying mitotic progression when chromosomes are incorrectly attached to the mitotic spindle. This delay is mediated by Mitotic Checkpoint Complexes (MCCs), which assemble at unattached kinetochores and repress the activity of the Anaphase Promoting Complex/Cyclosome (APC/C). The cellular localizations of MCCs are likely critical for proper SAC function, yet remain poorly defined. We recently demonstrated that in mammalian cells, in which the nuclear envelope disassembles during mitosis, MCCs diffuse throughout the spindle region and cytoplasm. Here, we employed binucleate yeast zygotes to examine the localization dynamics of SAC effectors required for MCC assembly and function in budding yeast, in which the nuclear envelope remains intact throughout mitosis. Our findings indicate that in yeast MCCs are confined to the nuclear compartment and excluded from the cytoplasm during mitosis. In contrast, we find that effectors of the Mitotic Exit Network (MEN) – a pathway that initiates disassembly of the anaphase spindle only when it is properly oriented – are in fact freely exchanged between multiple nuclei within a shared cytoplasm. Our study provides insight into how cell cycle checkpoints have evolved to function in diverse cellular contexts.

## Introduction

Accurate chromosome segregation during mitosis is facilitated by the Spindle Assembly Checkpoint (SAC), a conserved signaling pathway that monitors the attachment of chromosomes to the mitotic spindle via kinetochores, large protein complexes that assemble upon centromeric DNA^1–3^. Kinetochores form load-bearing attachments to spindle microtubules to facilitate: (1) chromosome alignment during prometaphase, and (2) segregation of sister chromatids during anaphase^3^. The SAC monitors kinetochore-microtubule attachment status, and delays anaphase onset in the presence of unattached kinetochores^4^, thus ensuring that when anaphase occurs, all chromosomes are positioned to be equally inherited. Through the activity of kinetochore-localized SAC effectors (*e.g*., Mad1, Mad2, Mad3, Bub1, and Bub3), unattached kinetochores generate a ‘wait anaphase’ signal, comprised of mitotic checkpoint complexes (MCCs)^5–8^. MCCs inhibit the activity of the Anaphase Promoting Complex/Cyclosome (APC/C), an E3 ubiquitin ligase, by sequestering the activating subunit Cdc20^9^. By inhibiting APC/C activity, MCCs prevent the degradation of key mitotic substrates such as Cyclin B and Securin, and thus delay anaphase onset^1^. In addition to Cdc20, the MCC is composed of Mad2, Mad3 (the homolog of human BubR1), and Bub3^9^. Mad1 and Bub1 catalyze the assembly of MCCs at unattached kinetochores, and are required for SAC function^10,11^.

Even a single unattached kinetochore is sufficient to delay anaphase onset^4,12^. Upon attachment of all kinetochores to spindle microtubules, MCC assembly ceases and cells rapidly enter anaphase^4,12,13^. The mechanisms that enable the SAC to maintain a robust mitotic delay, and yet also enable its rapid silencing remain unclear. One hypothesis explaining the robust nature of the checkpoint postulates that a single unattached kinetochore can catalyze the formation of sufficient levels of MCCs to maintain an arrest^14^. Cellular MCC concentrations are dictated by the rates of MCC assembly and disassembly, and the cellular volume that MCCs occupy during mitosis^14^. Alteration of these parameters perturb the strength of a SAC arrest^14–16^. For example, it is hypothesized that the high frequency of chromosome segregation errors observed in cells with large cytoplasmic volumes (*e.g*., embryonic cells and oocytes) results from the dilution of MCCs^15,17^. Our recent work characterized the mobility of MCCs within mammalian cells, and helped to define the relationship between cell volume and SAC activity^13^. Using fused mammalian cells (with two mitotic spindles)^18^, we demonstrated that both spindles in a fused cell entered anaphase synchronously, suggesting that MCCs can in fact move throughout the cytoplasm and between spindles. The parameters of MCC mobility in mammalian cells are dictated, in part, by the fact that these cells perform ‘open’ mitosis, in which the nuclear envelope breaks down upon entry into mitosis, thereby allowing mixing of the nucleoplasm and cytoplasm. This raises the question of how the presence of a nuclear envelope might impact the mobility of these effectors, and thus checkpoint function. Specifically, we wondered how nuclear-cytoplasmic exchange of SAC effectors – as affected by the nuclear import/export machinery – might impact SAC signaling. Moreover, how has the nuclear import/export machinery evolved to ensure proper SAC function in organisms in which the nuclear envelope remains intact (*i.e*., ‘closed’ mitosis)? We chose to use the budding yeast *Saccharomyces cerevisiae* to investigate these questions as these cells perform closed mitosis, and their SAC effectors are highly conserved with those found in metazoans. Here, we demonstrate that in contrast to mammalian cells, MCCs in yeast remain confined within the nucleus during mitosis, and that this confinement is likely important to ensure a robust checkpoint, which is important for high fidelity chromosome segregation.

## Results and Discussion

### Multiple Spindles Within a Shared Yeast Cytoplasm Initiate Anaphase Asynchronously

To determine if MCCs in budding yeast exchange between the nucleoplasm and cytoplasm, we generated binucleate yeast cells, and examined how two nuclear-confined mitotic spindles in a shared cytoplasm progress through mitosis. This approach has been used in both budding yeast and mammalian cells to determine the diffusible limits of cell cycle signals^13,18,19^. Attenuation of MCC assembly leads to activation of the APC/C, which in turn triggers anaphase onset^5–8^. Thus, a central assumption of these studies is that anaphase onset can only occur after MCC levels drop below the threshold required to inhibit APC/C activity. We therefore reasoned that if MCCs are restricted to the nucleoplasm, and not shared between nuclei within a binucleate cell, then the two spindles would enter anaphase independently of one another, thus exhibiting an asynchronous anaphase phenotype. Alternatively, if MCCs do in fact exchange between nucleoplasm and cytoplasm, then they would also be shared amongst the two nuclei. In this scenario, anaphase onset would be delayed until both spindles achieved proper kinetochore-microtubule attachments, at which point they would both initiate anaphase synchronously. We generated binucleate yeast cells by mating strains deleted for *PRM3* (*prm3*Δ), which is required for nuclear fusion during mating^20^. The resulting zygotes contained two nuclei in a shared cytoplasm (Fig. 1A). Mitotic spindles were visualized by GFP-Tub1 (α-tubulin) and Spc42-mCherry (a marker of spindle pole bodies, or SPBs; Fig. 1B). Newly formed zygotes were imaged every 2 minutes as they progressed through mitosis. Anaphase onset was defined as the time at which spindle elongation was initiated^21^.

**Figure 1.**
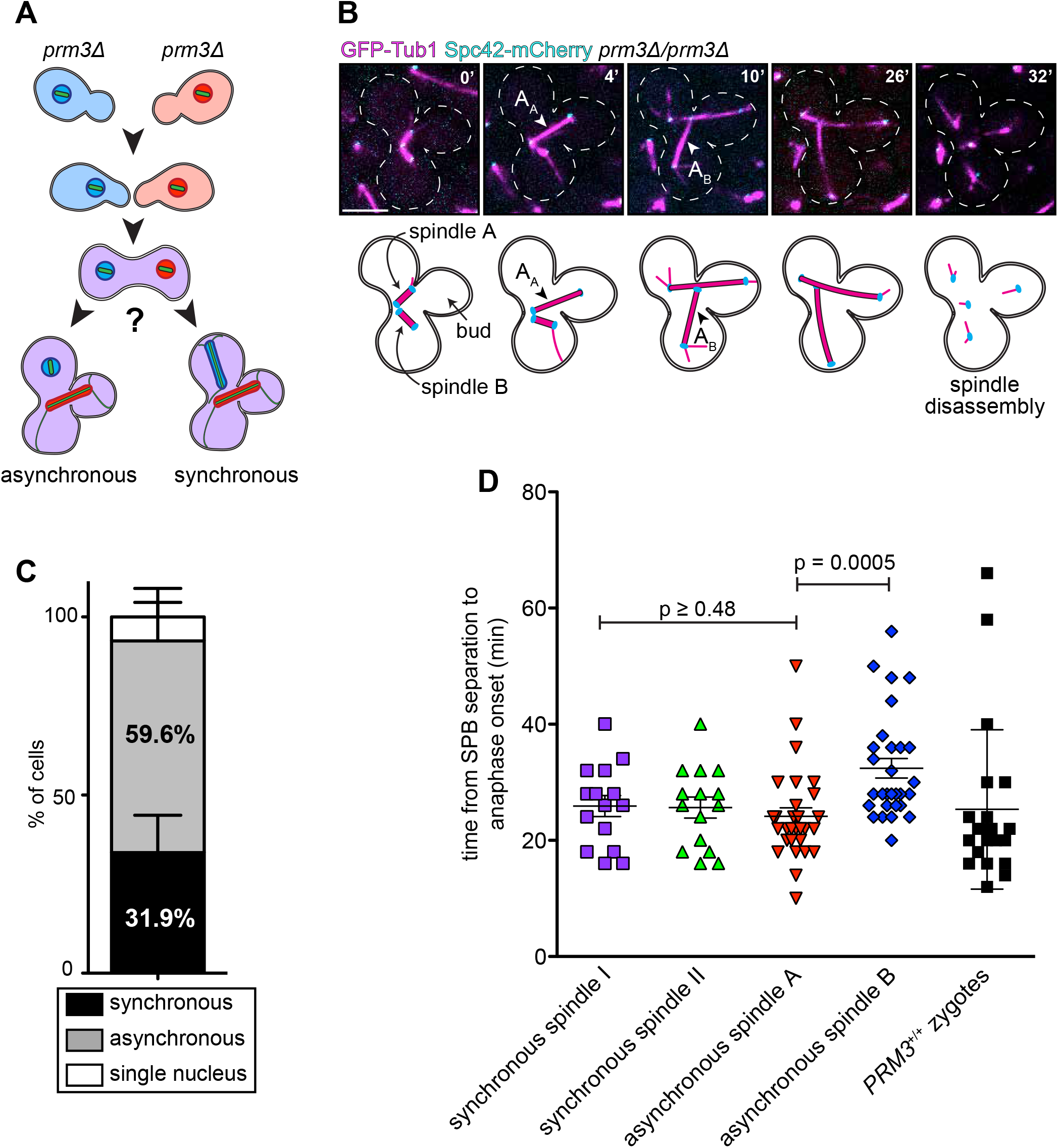
Anaphase onset occurs asynchronously in binucleate cells. (A) Schematic of experiment. *MATa prm3Δ* and *MATα prm3Δ* cells were mated together, generating binucleate zygotes. (B) Representative time-lapse images of asynchronous anaphase onset in a binucleate zygotes expressing GFP-Tub1 (magenta) and Spc42-mCherry (cyan). Spindle A initiates anaphase at 4’ (arrowhead, “A_A_”), while spindle B initiates anaphase at 10’ (arrowhead, “A_B_”). (C) Plot depicting the frequency of with which the indicated anaphase behaviors were observed (n = 47 binucleate zygotes from four separate experiments). Error bars denote standard deviation. (D) Plot depicting the duration of time spindles exhibiting the indicated anaphase onset behavior spent from spindle assembly (*i.e*., SPB separation) until anaphase onset. Zygotes were imaged every 2’ to determine the duration of mitosis. Error bars denote standard deviation. P values were calculated using two sided t-tests. Scale bars, 5 μm.

We found that the majority of cells (59.6%) displayed an asynchronous phenotype, in which anaphase onset for each spindle occurred at different times (Fig. 1B and C), whereas a minority (31.9%) exhibited synchronous anaphase onset. Figure 1B depicts representative time-lapse images in which the two spindles within a binucleate cell initiate anaphase at different times (*i.e*., are asynchronous; spindle A enters anaphase at 4’, “A_A_”, while spindle B enters anaphase at 10’, “A_B_”). Notably, both spindles disassemble at the same time (see below). In a small number of cells (8.5%), only one nucleus entered anaphase (Fig. 1C; “single nucleus”). These cells were excluded from further analysis.

To determine if the observed synchronous anaphase onset behavior was a consequence of one spindle waiting for the other (*i.e*., due to inter-nuclear MCC exchange), we measured the mitotic duration of spindles in synchronous and asynchronous cells. We predicted that if the first spindle to achieve metaphase – spindle I – waits for spindle II to also reach metaphase prior to synchronous anaphase onset, then spindle I would spend more time in mitosis. We defined mitotic duration as the time interval between spindle assembly (as determined by SPB separation) and anaphase onset. In contrast to our prediction, analysis of mitotic duration in cells exhibiting synchronous anaphase onset behavior revealed that each of the two spindles spent an equivalent amount of time in mitosis (Fig. 1D; p = 0.92). This value was also equivalent to the mitotic duration of asynchronous spindle A (the first spindle to initiate anaphase; p ≥ 0.48; Fig. 1D). Interestingly, we found that asynchronous spindle B (the second spindle to initiate anaphase) spent a significantly longer amount of time in mitosis than spindle A (Fig. 1D; p < 0.0005). Although the basis for the longer mitotic duration of B spindles is unclear, we hypothesize that it could simply represent stochastic differences in the efficiency of spindle assembly. Consistent with this notion, the range of mitotic duration values for A and B spindles (10-50 min, and 20-56 min, respectively) largely overlapped with the broad range of timing observed in wild-type (*PRM3*+/+) zygotes (Fig. 1D; 12-66 min). Furthermore, imaging of a chromosomally-integrated TetO array (using TetR-GFP) revealed that B spindles accurately segregated their chromosomes (Fig. S1). Specifically, we found that 100% of nuclei in zygotes exhibiting synchronous and asynchronous anaphase onset behavior accurately segregated chromosome IV (n = 8 synchronous nuclei, and 52 asynchronous nuclei). In summary, these findings indicate that synchronous anaphase events are coincident, and suggest that MCCs are confined to the nucleoplasm in yeast cells.

### Mad1 and Bub1 are Retained in the Nucleus Throughout the Cell Cycle

We next investigated the molecular mechanism by which individual nuclei within a shared cytoplasm exhibit autonomous SAC signaling. We hypothesized that nucleus-restricted localization of either individual checkpoint effectors, and/or assembled MCC complexes could explain this behavior. We first sought to determine the localization patterns of key SAC effectors throughout the cell cycle: Mad1, Mad2, Mad3, Bub1 and Cdc20 (Fig. 2A; also see Materials and Methods). In agreement with previous studies, we found that Mad1- and Mad2-GFP localized to the nuclear envelope throughout the cell cycle^22–25^, although the latter also exhibited diffuse localization in both the cytoplasm and nucleoplasm. Bub1-GFP localized as one or two foci per cell transiently during early mitosis, and was undetectable throughout the rest of the cell cycle. Previous studies have demonstrated that these Bub1-GFP foci likely coincide with kinetochores^26^. Finally, Mad3- and Cdc20-GFP exhibited diffuse cytoplasmic and nuclear localization that became enriched in the nucleus as cells progressed into mitosis, and rapidly decreased at the end of mitosis.

**Figure 2.**
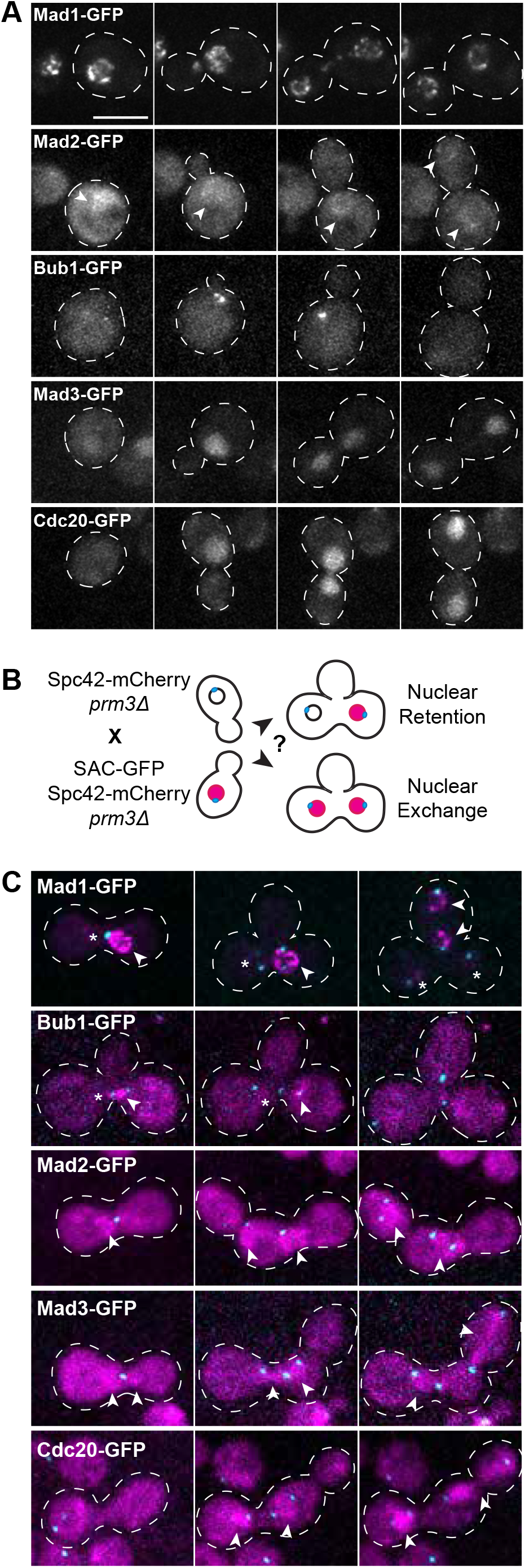
SAC effectors exhibit variable localization dynamics throughout the cell cycle. (A) Representative time-lapse images of haploid cells expressing Mad1-, Mad2-, Bub1-, Mad3-, or Cdc20-GFP as they progress through mitosis. Arrowheads in Mad2-GFP panel denote nuclear envelope localization. (B) Schematic depicting experimental approach to determine the localization dynamics of test SAC effectors in binucleate zygotes. (C) Representative time-lapse images of binucleate zygotes expressing denoted test SAC-GFP. Arrowheads denote nuclear GFP fluorescence, and asterisks denote nuclei that lack GFP signal. Images were acquired every 5’. Scale bars, 5 μm.

We next asked whether any of these factors were restricted to individual nuclei within binucleate cells. To address this, we mated *prm3*Δ cells expressing a GFP-tagged SAC effector to *prm3*Δ cells that did not express the GFP-fusion (Fig. 2B and C). To delineate the approximate nuclear position, these cells also expressed Spc42-mCherry, which marks the nuclear envelope-embedded SPBs. We also attempted to delineate the nuclear envelope or nuclear compartment using mCherry tagged alleles of the nucleoporin Nup133 or the histone Htb2 (data not shown). However, the fluorescent signal generated by both Nup133-mCherry and Htb2-mCherry was detected in the GFP channel, and confounded our analysis of GFP-tagged SAC effector exchange. We imaged cells from the time of fusion until anaphase onset, and assessed to what extent, if any, the GFP-SAC effector localized to both nuclei. Interestingly, we found that both Mad1- and Bub1-GFP remained exclusively enriched in 100% of GFP-expressing nuclei, and were never observed in the other nucleus (Fig. 2C; non-SAC-GFP-expressing nucleus marked with an asterisk; n ≥ 10 cells). This suggests that these proteins – which are both catalysts of MCC assembly – remain confined within the nucleus throughout the cell cycle, and are not shuttled into the cytoplasm or exchanged between nuclei. In contrast, both nuclei in binucleate zygotes accumulated the MCC complex components Mad2-, Mad3-, and Cdc20-GFP prior to spindle assembly (as assessed by SPB separation; see arrowheads in Fig. 2C; n ≥ 10 cells), suggesting that these factors are indiscriminately imported into both nuclei early in the cell cycle. These findings indicate that checkpoint effectors in yeast exhibit distinct localizations throughout the cell cycle, and also exhibit different nuclear import/export properties.

### Cdc20 Is Retained in the Nucleus Subsequent to Import

Given that neither Mad1 nor Bub1 – both of which are nuclear confined – are MCC subunits^9,11,27^, the mechanism by which they could provide nuclear autonomy with respect to anaphase onset is unclear. Our observations indicate that Mad2, Mad3 and Cdc20 – which are catalytically assembled into the MCC at unattached kinetochores^9,11,27^ – are indiscriminately imported into both nuclei early in the cell cycle. Yet, the asynchronous anaphase onset behavior suggests that intact MCCs are confined to the nucleoplasm. Thus, we hypothesized that subsequent to nuclear import, MCC subunits or intact complexes become confined within the nucleus. To determine if this is the case, we used a strategy that would permit us to quantitatively determine the degree of nuclear exchange of Cdc20 subsequent to its import. To this end, we employed combined FLIP (Fluorescence Loss In Photobleaching)-FRAP (Fluorescence Recovery After Photobleaching) analysis of Cdc20-GFP-expressing binucleate cells.

As proof-of-concept, we performed FLIP-FRAP with Arx1, a known nuclear shuttling factor that is similar in molecular weight to Cdc20^28^, and thus likely exhibits similar passive nuclear import/export parameters. Studies have shown that proteins smaller than 50 kDa can passively diffuse through the nuclear pore, while proteins larger than 50 kDa rely on karyopherin-mediated import and export^29,30^. Both Arx1 and Cdc20 are greater than 50 kDa (65 and 67 kDa respectively; 94 kDa and 96kDa with GFP), and thus likely require active transport to transit through nuclear pores.

When we photobleached a single Arx1-GFP-containing nucleus (Fig. 3; “nucleus 1”) in a binucleate cell, the GFP fluorescence recovered to 43.0% of its original value after two minutes (after correcting for photobleaching; see Materials and Methods), which is due to the import of unbleached Arx1 into this nucleus. Conversely, the fluorescence intensity of the unbleached Arx1-GFP-containing nucleus (“nucleus 2”) decreased by 25.3% after two minutes, indicating that Arx1 molecules from nucleus 2 were actively exported over this time frame. These data support the notion that Arx1 is indeed exchanged between nucleoplasm and cytoplasm^28^.

**Figure 3.**
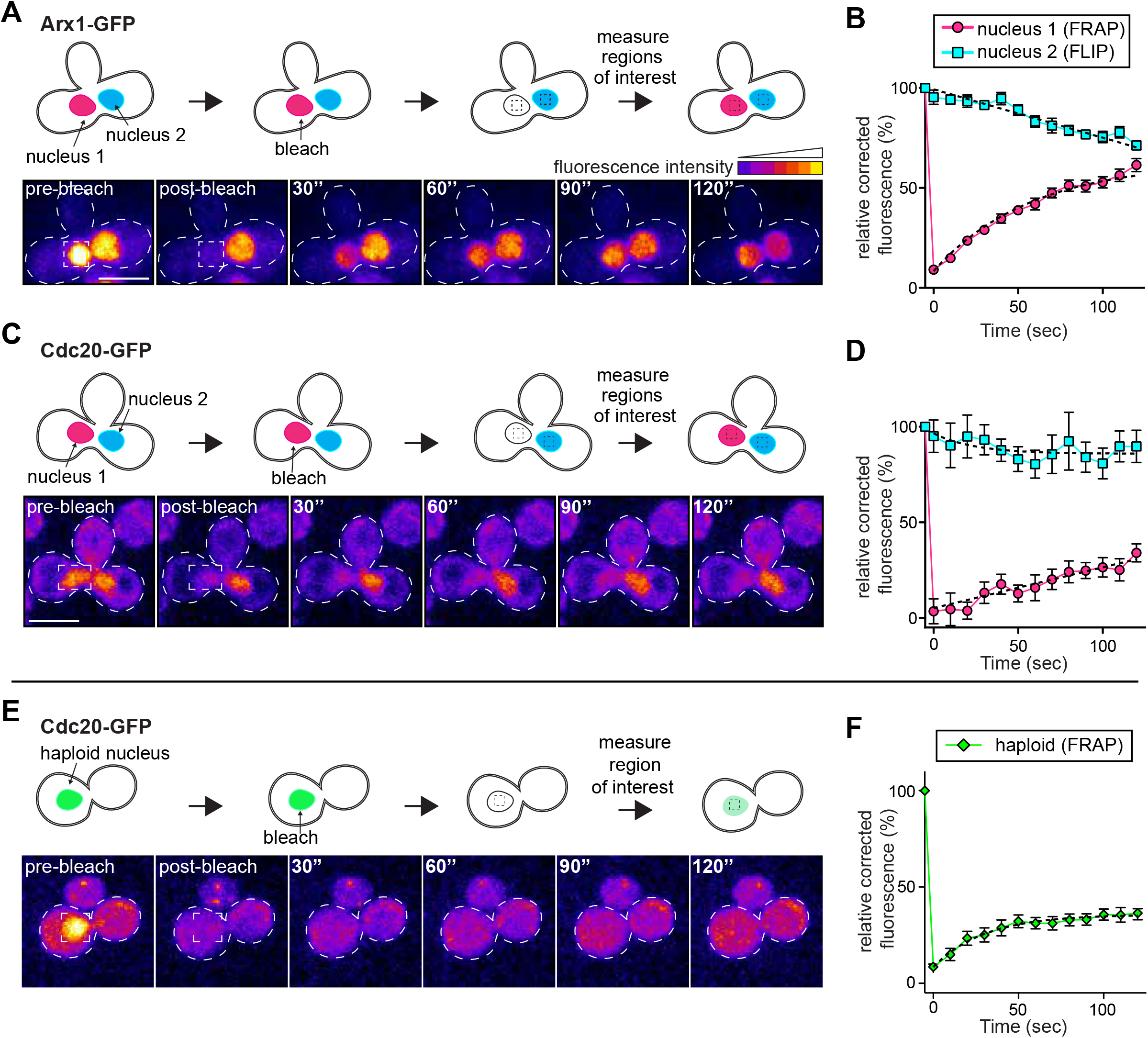
Cdc20 is restricted to the nucleus during mitosis. (A, C and E) Schematic of experimental setup along with representative time-lapse images depicting FRAP-FLIP analysis of (A) Arx1-GFP-, or (C) Cdc20-GFP-expressing binucleate zygotes, or (E) Cdc20-GFP-expressing haploid cells. Fluorescence intensities are displayed as a heat map. (B and D) Relative corrected average fluorescence recovery in the photobleached nucleus (nucleus 1, FRAP; magenta circles), or loss of fluorescence in the unbleached nucleus (nucleus 2, FLIP; blue squares) plotted over time for (B) Arx1- and (D) Cdc20-GFP-expressing zygotes (n ≥ 10 binucleate zygotes; see Materials and Methods). (F) Relative corrected average fluorescence recovery in the photobleached nucleus of Cdc20-GFP-expressing haploid cells (green diamonds; n ≥ 10 cells). Error bars denote standard error. Curve fits (dashed lines) are one-phase decay non-linear regressions fit to the experimental data. Images were acquired every 10’’ for 120’’. Scale bars, 5 μm.

In contrast to Arx1, FRAP analysis of Cdc20-GFP revealed a lesser extent of fluorescence recovery in nucleus 1 after two minutes of recovery (27.9%; Fig. 3C and D). More strikingly, FLIP analysis of nucleus 2 revealed almost no loss of fluorescence (4.8%). In light of the minimal fluorescence loss in nucleus 2, we hypothesized that the 27.9% fluorescence recovery of Cdc20-GFP in nucleus 1 was due to the import of Cdc20 molecules from the cytoplasm. To test this, we performed FLIP-FRAP experiments on Cdc20-GFP expressing haploid cells, in which nuclear fluorescence recovery can only be due to the import of unbleached proteins from the cytoplasm. This analysis revealed that the same degree of fluorescence recovery occurred in these cells as in binucleate zygotes (27.9% after two minutes for both; Fig. 3F). Thus, the nuclear fluorescence recovery of Cdc20-GFP in binucleate cells is likely due to import of protein from the cytoplasm, and not to exchange between nuclei. Taken together, these results suggest that upon import into the nucleus, Cdc20 is restricted from export back to the cytoplasm. Due to the poor signal-to-noise ratio, we were unable to perform similar experiments on other MCC components (*e.g*., Mad2 and Mad3). However, given the observed asynchrony of anaphase onset in 59.6% of binucleate cells, and our FLIP-FRAP data for Cdc20 – a key component and substrate of the MCC – we postulate that upon nuclear import and subsequent assembly of Cdc20, Mad2, Mad3 and Bub3 into intact MCCs, these complexes are nuclear confined. We propose that the asynchronous anaphase onset behavior observed in binucleate zygotes is a direct consequence of the nuclear confinement of intact MCCs.

Although the mechanisms underlying nuclear confinement of MCCs in this organism are unclear, it stands to reason that SAC activity could be impacted, at least in part, by nuclear import/export-dependent processes. A growing body of work indicates that nuclear import/export is subject to cell cycle specific regulation, and is important for cell cycle transitions^22,23,31^. The molecular basis of MCC nuclear confinement, and the relevance of this sequestration to overall SAC function in yeast will be an important future direction of study. Given that SAC strength correlates with the volume which MCCs occupy^15,17^, one possibility is that nuclear sequestration of MCCs ensures their appropriate nuclear concentrations, thus enabling proper SAC function. Thus, we predict that shifting the localization dynamics of the MCC to favor increased nuclear export may weaken the SAC, and thus lead to an increased prevalence of chromosome segregation errors.

### Mitotic Exit Occurs Synchronously in Binucleate Zygotes

While studying the anaphase onset behavior in binucleate cells, we observed that in 83.3% of cells, disassembly of the two mitotic spindles occurred simultaneously (Fig. 1B, and Fig. 4C and D). Spindle disassembly is regulated by the Mitotic Exit Network (MEN), a signaling pathway that ensures that spindle disassembly occurs only when the spindle is properly oriented through the bud neck^32,33^. It is well established that the key effector of MEN signaling, Tem1, resides at SPBs^34^. When both SPBs of an anaphase spindle are situated within the mother cell compartment (as in the case of a mispositioned spindle), MEN inactivity ensures that spindle disassembly and consequent mitotic exit do not occur^35,36^. However, upon entry of one SPB into the bud, the MEN is activated and mitotic exit ensues^36^. A recent study using binucleate zygotes demonstrated that entry of one SPB into the bud (from one mitotic spindle) was sufficient to activate the MEN, even if both SPBs from the other spindle were situated within the mother cell^37^. From this, the authors concluded that mispositioned spindles do not generate a dominant inhibitory signal, but rather a properly positioned spindle generates an activating signal that enables the MEN to promote mitotic exit.

**Figure 4.**
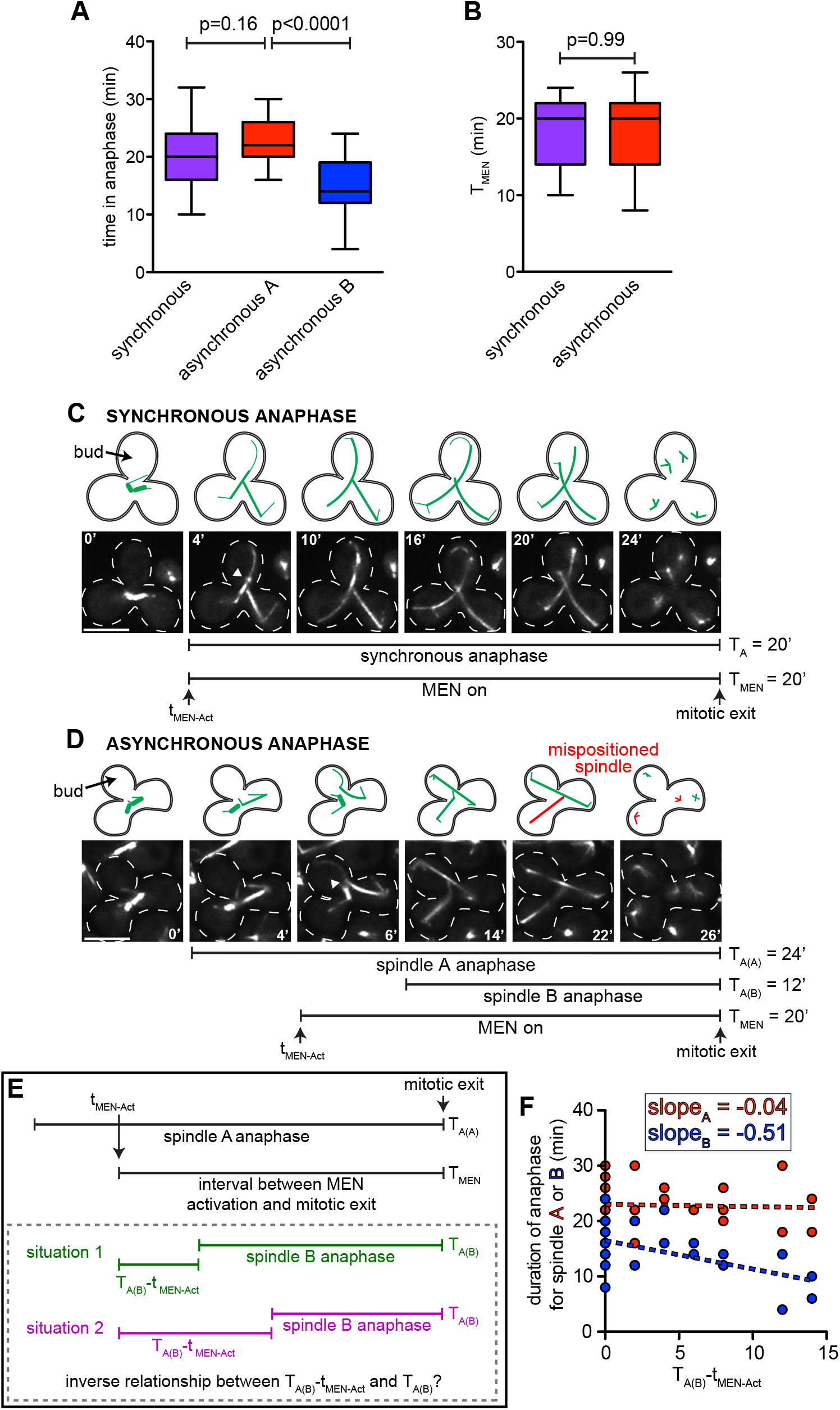
Mitotic exit occurs synchronously in binucleate zygotes. (A) Box-whisker plot depicting the duration of anaphase for spindles exhibiting indicated anaphase onset behaviors (n ≥ 24 cells). (B) Box-whisker plot depicting the duration of time over which the MEN is active (*i.e*., the time between entry of the first SPB of an anaphase spindle into the bud, and spindle disassembly, or mitotic exit, T_MEN_) for spindles exhibiting the indicated anaphase onset behavior (n ≥ 25 zygotes). (C) Representative time-lapse images, along with cartoon model, of anaphase onset and MEN behavior in a zygote exhibiting synchronous anaphase onset (T_A_, duration of anaphase). Note that both spindles enter anaphase, and the SPB of one spindle enters the bud at 4’ (arrowhead). 20 minutes later, both spindles disassemble, and the cell exits mitosis. (D) Representative time-lapse images, along with cartoon model, of anaphase onset and MEN behavior in a zygote exhibiting asynchronous anaphase onset (T_A(A)_, duration of anaphase for spindle A; T_A(B)_, duration of anaphase for spindle B). (E) Predicted correlation between T_A(A)_ or T_A(B)_ and the time between anaphase onset of spindle B (T_A(B)_) and MEN activation (tMEN-Act). (F) Plot depicting relationship between T_A(A)_ (red circles) or T_A(B)_ (blue circles) and TA(B)-tMEN-Act (n = 24). Dashed lines represent linear regressions modeled from raw data, along with slopes. Data points that fall at T_A(B)_-t_MEN-Act_ = 0 represent instances when the B spindle entered anaphase at the same time as the MEN was activated. Images were acquired every 2’. Scale bars, 5 μm. For box-whisker plots, whiskers define the range, boxes encompass 25^th^ to 75^th^ quartiles, and lines depict the medians.

In agreement with this model, we also found that mitotic exit occurred in the presence of mispositioned spindles (Figs. 1B and 4D). To confirm that a single anaphase spindle could indeed generate a dominant MEN activating signal (*i.e*., sufficient to cause mitotic exit in both spindles within the cell), and to gain further insight into this process, we performed an in-depth analysis of anaphase behavior in these cells. Since asynchronous spindles A and B initiated anaphase at different times, yet disassembled at the same time, we predicted that either their rates of elongation would differ, or that they would achieve different maximum lengths prior to disassembly. Consistent with our prediction, we found that although asynchronous spindles A and B elongated at nearly identical rates (and matched the rate of spindle elongation in synchronous cells; Fig. S2A), spindle A did in fact achieve a significantly longer maximum length than spindle B (Fig. S2B; p = 0.04). Moreover, we found that on average, asynchronous spindle B spent 52.4% less time in anaphase than spindle A (Fig. 4A; p < 0.0001), the latter of which did not differ significantly from our measurement of spindles that underwent synchronous anaphase (p = 0.16). These data suggest that spindle B exits mitosis prematurely.

We wondered whether activation of the MEN by spindle A is responsible for the premature disassembly of spindle B. To test this, we measured the duration of time between MEN activation (*i.e*., the point at which the first SPB of an anaphase spindle crosses the bud neck, tMEN-Act) and mitotic exit (*i.e*., spindle disassembly) in binucleate zygotes (T_MEN_; see Fig. 4E). We found that the T_MEN_ values in zygotes undergoing synchronous and asynchronous anaphase onset were not significantly different (p = 0.99; Fig. 4B-D). This suggests that regardless of the state of mitotic progression of spindle B, spindle A-mediated activation of the MEN will result in cell-wide mitotic exit after a fixed duration (T_MEN_ ~ 18.5 minutes; Fig. 4B). If true, we reasoned that the longer it takes for spindle B to enter anaphase relative to the time of MEN activation, the less time it will spend in anaphase (see Fig. 4E; T_A(B)_). Moreover, if spindle A is responsible for generating the dominant MEN activation signal – which occurs independently of spindle B behavior – then there should be no such relationship for spindle A. To determine if this was the case, we correlated the duration of anaphase for spindles A or B with the time interval between MEN activation and anaphase onset of spindle B (T_A(B)_-t_MEN-Act_; Fig. 4E and F). We found that there was indeed an inverse relationship between T_A(B)_-t_MEN-Act_ and the duration of anaphase for spindle B; however, no such correlation was observed for spindle A (Fig. 4F). Taken together, these observations confirm the notion that, in contrast to the nuclear-confined SAC machinery, the signal that activates the MEN is dominant and can trigger a cell-wide response.

### Materials and Methods

#### Strain generation, culture methods, and preparation for imaging

All yeast strains were constructed in the BY4743^38^ background and are listed in Table S1. TetO/TetR-GFP strains were gifts from Dr. Jeffrey Moore (University of Colorado, Anschutz Medical Campus), and those expressing GFP-SAC effectors were gifts from Dr. Santiago DiPietro (Colorado State University). It is worth noting that some of the GFP-tagged alleles used in this study, such as Mad2, are hypomorphic. Because we did not rely on the checkpoint function of these proteins for data interpretation, we did not assess the functionality of these alleles using standard benomyl sensitivity assays. Cells were grown and maintained in rich (YPD) or synthetic defined (SD) media at 30°C^39^. Transformations were performed using the standard lithium acetate method. Strains expressing fluorescently tagged proteins were gifts (see above) or constructed by either PCR product transformation, plasmid integration, or by mating and tetrad dissection. To mate cells for generation of zygotes, parental strains were grown in YPD overnight at 30°C. The next day, approximately equivalent numbers of cells were mixed together in 50 μL of YPD, spotted on a YPD plate, and incubated at 30°C for 3-4 hours. Cells were then scraped from YPD plate, washed twice with SD media, and prepared for imaging.

#### Plasmid generation

Plasmids used in this study are listed in Table S2. To produce cells with fluorescently-labeled spindle pole bodies, we generated a plasmid that would integrate mCherry::HYG^R^ (encoding hygromycin resistance) at the 3’ end of the *SPC42* locus. To this end, the 3’ end of *SPC42* (nucleotides 699-1089) was PCR amplified using primers flanked with ClaI restriction sites on the 5’ and 3’ ends, digested with ClaI, and ligated into *pmCherry::HYG^R^* digested similarly. This plasmid, pSPC42-mCherry:: HYG^R^, was digested with AflII prior to transformation and selection on hygromycin-containing media.

#### Live cell microscopy

All microscopy was carried out on an inverted Nikon Ti-E microscope equipped with a Perfect Focus unit, a 1.49 NA 100X CFI Plan Apo objective, a piezoelectric stage (for Z-control), an electronically controlled emission filter wheel, an iXon X3 DU888 EM CCD camera (Andor), and a Yokagawa spinning disc head. Excitation light (for imaging and targeted photobleaching) was provided by an AOTF-controlled laser launch with 7 lines (Nikon; 405 nm, 445 nm, 488 nm, 514 nm, 561 nm, 594 nm, 640 nm), and two outputs (one dedicated to the spinning disk head, and the other to a PA/FRAP unit). The system was controlled by NIS-Elements running on a 64-bit workstation. For time-lapse imaging, cells were perfused into a CellASIC ONIX microfluidics chamber (plate type Y04C, for haploid yeast cells; Millipore). SD media was continuously perfused into the imaging chamber at 7 psi, and the chamber was maintained at 30°C throughout the experiment. Step sizes of 0.5 μm were used to acquire 3.5 μm thick Z-stacks every 2, 2.5, or 5 minutes (as indicated in figure legends). For FRAP-FLIP (see below), cells were spotted onto a 1.7% SD agarose pad. After ~1 minute, a coverslip was mounted on top of the cells, and sealed with paraffin wax.

#### Image analysis and processing

Time-lapse images were analyzed in both NIS-Elements and ImageJ Fiji (ImageJ, National Institutes of Health) programs. Mitotic spindle lengths were measured and calculated in 3-dimensions. All images presented throughout this study are maximum intensity Z-projections. All brightness and contrast modifications were performed in Adobe Photoshop. Heat-map intensity images presented in Figure 3 were prepared in ImageJ Fiji after images had been modified in Adobe Photoshop (identical brightness/contrast settings were used for all images within a given experiment). All statistical analyses were performed using GraphPad Prism. Significance was calculated using unpaired t-tests.

#### Fluorescence Recovery After Photobleaching (FRAP) and Fluorescence Loss In Photobleaching (FLIP)

Photobleaching was performed using a 20 mW 405 nm laser at 25% power. After acquisition of a pre-bleach image (exposures: 200 ms, Arx1-GFP; 300ms Cdc20-GFP), a single focused 25 ms laser pulse was used to photobleach one nucleus in a binucleate cell. The pulse reduced GFP fluorescence by 70-95%. The extent of fluorescence reduction following the pulse was taken into account when calculating the degree of recovery (43.0% and 27.9% for Arx1- and Cdc20-GFP, respectively). Immediately following the targeted bleach, 0.5 μm step sizes were used to acquire 1.5 μm thick Z-stacks every 10 seconds for 120 seconds. Control cells (n ≥ 5 cells for both Cdc20-GFP and Arx1-GFP experiments) were subjected to an identical imaging sequence, but without the targeted photobleach pulse. To correct for non-targeted photobleaching, the calculated fluorescence loss in control cells was fitted to a linear regression in Graphpad Prism. The signal loss calculated from the regression equation at each time point was added to both the calculated FRAP and FLIP experimental values. Using ImageJ, the mean fluorescence intensity values for a 5×5 pixel region of interest in each nucleus were corrected for background fluorescence and photobleaching during image acquisition and plotted as the mean intensity with standard error. Graphpad Prism software was used to fit these data to single-decay non-linear regressions.

## Acknowledgments

We would like to thank Dr. Santiago DiPietro (Colorado State University) and Dr. Jeffrey Moore (University of Colorado, Anschutz Medical Campus) for sharing yeast strains. This work was supported by the National Institutes of Health (R01 GM088731 to J.G.D. and R01 GM118492 to S.M.M.) and the National Science Foundation (1518083 to J.G.D. and S.M.M.). L.R.H. designed and executed the study, and helped write the manuscript. S.M.M. helped write the manuscript, and S.M.M. and J.G.D. provided funding, and helped to edit the manuscript.

## Abbreviations

SAC: Spindle Assembly Checkpoint
MCC: Mitotic Checkpoint Complex
APC/C: Anaphase Promoting Complex/Cyclosome
FLIP: Fluorescence Loss In Photobleaching
FRAP: Fluorescence Recovery After Photobleaching
GFP: Green Fluorescent Protein
SPB: Spindle Pole Body

